# Seed predation increases from the Arctic to the Equator and from high to low elevations

**DOI:** 10.1101/304634

**Authors:** A.L. Hargreaves, Esteban Suárez, Klaus Mehltreter, Isla Myers-Smith, Sula E. Vanderplank, Heather L. Slinn, Yalma L. Vargas-Rodriguez, Sybille Haeussler, Santiago David, Jenny Muñoz, Roberto Carlos Almazán Núñez, Deirdre Loughnan, John W. Benning, David A. Moeller, Jedediah F. Brodie, Haydn J.D. Thomas, M.P.A. Morales

## Abstract

Species interactions have long been predicted to increase in intensity towards the tropics and low elevations, due to gradients in climate, productivity, or biodiversity. Despite their importance for understanding global ecological and evolutionary processes, plant-animal interaction gradients are particularly difficult to test systematically across large geographic gradients, and evidence from smaller, disparate studies is inconclusive. By systematically measuring post-dispersal seed predation using 6980 standardized seed depots along 18 mountains in the Pacific cordillera, we found that seed predation increases 18% from the Arctic to Equator and 16% from 4000 masl to sea level. Clines in total predation, likely driven by invertebrates, were consistent across tree-line ecotones and in continuous forest, and were better explained by climate seasonality than by productivity, biodiversity, or latitude. These results suggest that species interactions play predictably greater ecological and evolutionary roles in tropical, lowland, and other less seasonal ecosystems.

**One Sentence Summary:** Post-dispersal seed predation increases from the Arctic to the Equator and from high elevations to sea level.

---

Few biological patterns are as striking as latitudinal and elevational changes in biotic communities. Biodiversity and ecosystem productivity increase dramatically toward low latitudes (1, 2) and elevations (3, 4). Biologists have long speculated that greater diversity and productivity should generate corresponding increases in the intensity of species interactions (5-7). However, tests for gradients in interaction intensity (8-12) or their expected ecological and evolutionary signatures (e.g. density dependence 13, 14, defenses 15, 16) find contradictory results. While latitude and elevation are often considered analogues, their effects on interaction strength are rarely tested together. This likely contributes to the variability of experimental results, and limits our understanding of their joint effects on global patterns in species interactions.

More intense interactions toward low latitudes and elevations underpin several iconic biogeographic hypotheses. Antagonistic species interactions are thought to maintain high tropical diversity by limiting species dominance (the Janzen-Connell hypothesis; 17, 18), amplify tropical diversity by accelerating speciation (7, 19), and play a predictably greater role in determining species’ warm (low-latitude and elevation) vs. cool range limits (5, 6). For example, stronger tropical seed predation—an interaction that shapes plant communities and distributions (20, 21)—is proposed to explain the greater tropical diversity of trees (14, 17, 18) and adaptations for seed defense (22). The strength and predictability of interaction gradients is therefore pivotal to understanding their role as macroevolutionary and biogeographic agents.

Despite an outsized role in theory, assessing the generality of interaction gradients is hampered by constraints of existing evidence (23). Most studies encompass a limited spatial scale, omitting one or both latitudinal extremes (tropics or tundra; 11, 12, 24). Particularly lacking are replicated elevational gradients across a significant latitudinal range. Another challenge is controlling for evolutionary responses that can mask underlying gradients, such as increased plant defenses to counter chronically high consumption (11, 15). Finally, gradients in interaction strength can differ between taxa (e.g. invertebrates vs. vertebrates), counteracting each other’s effects when combined (8, 9). Notably, large-scale experiments using standardized methods and materials have generated the most convincing examples of species interaction gradients (9, 25, 26). Here, we expand on previous work by explicitly testing both latitudinal and elevational gradients in a globally-important plant-animal interaction—post-dispersal seed predation.

While the generality of geographic patterns in interactions has been extensively debated, their causes have received less attention (9, 23). Coarse gradients could arise if seed predation differs among biomes (e.g. declines above treeline (27)), and low-predation biomes are more common at high latitudes and elevations. This could explain why studies in single biomes sometimes fail to find latitudinal patterns (12, 24). Alternatively, seed predation could increase with factors that continuously increase granivore populations or feeding rates towards tropical and lowland areas (28). Warmer, less variable temperatures can increase animal populations and activity, especially for invertebrates (29, 30). Greater vegetation productivity could increase granivore populations by enhancing available food and niche space (24). Though rarely tested, higher diversity of seed predator communities could increase total granivore numbers by minimizing competition and maximizing resource use (31, 32).

In line with theory, we predicted that post-dispersal seed predation would become more intense from high latitudes to the Equator and from high elevations to sea level. We expected invertebrate seed predation to show particularly strong patterns in relation to geography (latitude and elevation) and climate (biome and temperature), as temperature directly affects ectotherm activity (29, 30). If invertebrates drive interaction gradients (as in 9), we expect total and invertebrate seed predation to show similar gradients—i.e. vertebrates contribute a consistent amount of predation. We predict invertebrate-driven seed predation to decline both sharply above treeline, as major invertebrate groups ‘drop out’ (27, 33), and continuously with falling temperature due to metabolic constraints.

We tested our predictions by systematically measuring predation on 6980 groups (‘depots’) of standardized seeds at 79 sites from 25 to 4120 meters above sea level (masl) and from 0° to 64°N. Sites were arranged in 18 elevational transects along the west coast of the Americas. Each transect comprised 4-5 sites covering a median of 1007 m elevation (Fig. 1A, Fig. S1). We used one oil-based and one starch-based agricultural seed species: sunflower (*Helianthus annuus*) seeds without shells, and oat (*Avena sativa*) seeds with husks. Seeds were bulk-purchased from a consistent supplier and heat treated to make them inviable. We ran the experiment 56 times from 2015-2017, with each transect tested 1-6 times (median=4). Each run of the experiment, we placed 20 depots of 8 sunflower seeds and 10 depots of 5 oat seeds at each site along a transect, with depots alternating between seed types and separated by ≥5 m. To isolate invertebrate seed predation, we secured wire vertebrate-exclusion cages over 3 or 4 sunflower depots per site during 25 experimental runs (60 sites; Fig. S2). After 24 h we counted the seeds that were intact, partially eaten, or missing. For each 24 h assay, we calculated the mean fraction of seeds predated ((eaten+missing)/initial; 70-99% of seeds removed from the ground are eaten 21, 34) for each seed type and predator type (total or invertebrate) at each site. We used generalized linear mixed-effects models (GLMMs; 35) to quantify the effect of latitude, elevation, seed type, vegetation biome, and their interactions on seed predation. We used structural equation models to compare the explanatory power of climate (temperature and precipitation mean and annual variation), actual evapotranspiration (AET, a proxy for ecosystem productivity), and vertebrate richness (a proxy for granivore diversity), downloaded at 1-10 km^2^ resolution for each site (35).

Consistent with long-standing predictions, seed predation increased toward the tropics and low elevations (Fig. 1, full GLMM results are in Table S2). Total seed predation was 10% higher in the tropics than in the temperate zone (south vs. north the Tropic of Cancer: likelihood ratio test χ^2^_df=1_ 20.7, *P* < 0.001, Fig. 1G) for oil- and starch-based seeds (geographic patterns did not vary between sunflower and oat seeds in any analysis, Fig S3). Seed predation increased by 2.8% for every 10° of latitude closer to the equator (latitude: χ^2^_df=1_ 17.7, *P* < 0.001; Fig. 1B), and by 0.4% for every 100 m decline in elevation (elevation: χ^2^_df=1_ 6.92, *P* = 0.009), independent of latitude (elevation × latitude interaction: χ^2^_df=1_ 2.54, *P* = 0.11; Fig. 1C). In total, seed predation increased by 18% from Alaska to Ecuador and 16% from 4000 masl to sea level—important experimental evidence that interactions have a greater potential to shape tropical and lowland communities.

**Fig. 1.**
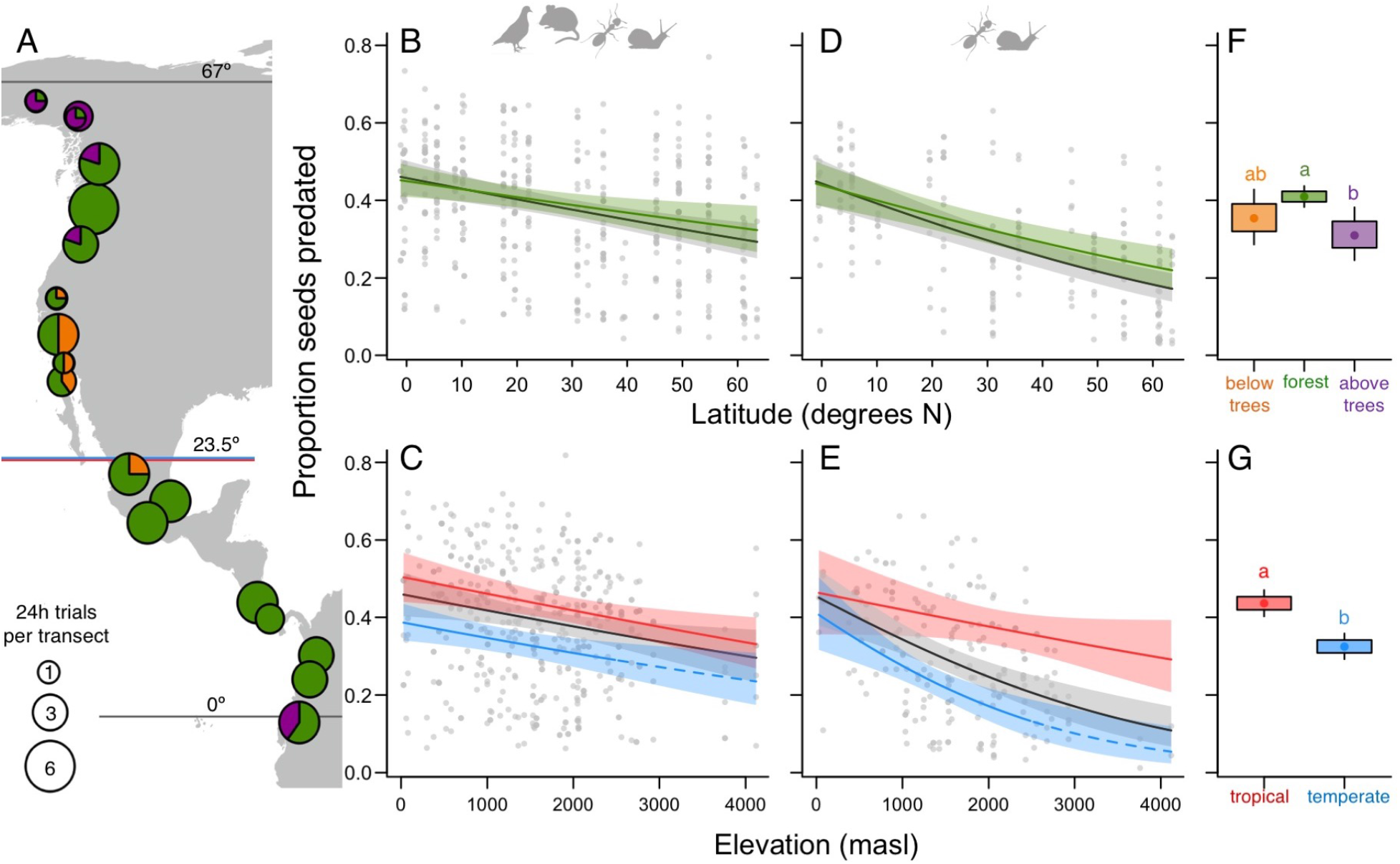
Latitudinal and elevational declines in seed predation. (A) Sampling transects from the Arctic to Equator. Circle area is proportional to the mean times the experiment was run at each site on the transect (1 to 6), pie slices show the proportion of sites per biome: above upper tree line, forest, below lower tree line. (B-E) Geographic trends in predation on sunflower seeds (+/-95% confidence intervals) fitted by generalized linear mixed models – patterns did not differ for oat seeds (35). Latitudinal trends (B, D) are shown for 1500 m (median) elevation across biomes (black lines) and in forests (green lines). Elevational trends (C, E) are shown for the median latitude (31°N; black lines), tropics (<23.5°N; red lines), and temperate zone (>23.5°N; blue lines). Dashed trend lines show model extrapolations for temperate sites above 2500 m. Latitudinal and elevational patterns are similar for total seed predation (B, C: 56 experimental runs) and predation by invertebrates (D, E: 25 experimental runs). Points show partial residuals for the all-site model (black) in each panel. Biomes (F) and latitudinal zones (G) differed significantly in total seed predation (different letters indicate significant differences) and invertebrate seed predation (not shown). Dots, boxes and lines show the mean, 1 SE, and 95% confidence interval, respectively. Full statistical results in Table S2.

Latitudinal and elevational clines in invertebrate seed predation paralleled clines in total predation, suggesting geographic patterns are driven primarily by invertebrates. For the subset of sites and dates where we excluded vertebrates from some depots, elevational gradients in seed predation were steeper at high vs. low latitudes (latitude × elevation: χ^2^_df=1_ 7.5, *P* < 0.01; Fig. 1E), but patterns were consistent between total and invertebrate predation (no interactions between vertebrate-exclusion treatment and other variables: χ^2^_df=3_ 2.1, *P* = 0.55). Parallel gradients in total vs. invertebrate seed predation suggest that vertebrate seed predation is relatively constant, meaning invertebrates mediate increased predation toward low latitudes and elevations.

Both categorical differences among biomes and continuous environmental gradients contributed to gradients in seed predation. Even after accounting for elevation, seeds above treeline experienced the lowest total predation (biome: χ^2^_df=1_ 8.0, *P* = 0.019; Fig. 1F) and invertebrate predation (biome, vertebrate-exclusion depots only: χ^2^_df=1_ 8.1, *P* = 0.017). However, when we restricted analyses to forests (72% of our sites), patterns in seed predation were weaker— demonstrating that biome differences contribute to interaction patterns, but persisted— demonstrating that continuous ecological gradients contribute as well (Fig. 1B, D), with particularly strong effects of elevation at high latitudes (elevation × latitude, total predation: χ^2^_df=1_ 4.2, *P* = 0.041; invertebrate predation: χ^2^_df=1_ 3.7, *P* = 0.05).

Among the continuous ecological factors examined, annual temperature range explained total seed predation better than other climate variables, productivity (AET), or diversity, particularly for sunflower seeds (model with annual temperature range + elevation: R^2^ = 0.366, Table S3). Total seed predation increased as temperature range declined and environments became less seasonal (Table S4). Contrary to our prediction of stronger temperature effects on invertebrates, latitude explained invertebrate seed predation better than more mechanistic predictors, including climate (Tables S3-4). Our results add to mounting evidence relating biotic interaction strength to climate (24, 36, 37), though we found a less direct role of temperature and a stronger role for seasonality than expected.

Overall, our study reveals strong latitudinal and elevational gradients in the intensity of species interactions. Latitudinal and elevational patterns held for oil- and starch-based seed, total and invertebrate predation, across biomes and within forests. Large-scale patterns emerged despite variation among elevational gradients (Fig. S1) and even between replicated experiments on the same gradient—variation that has probably contributed to equivocal results from studies at smaller spatial scales and with less temporal replication (8). Our results are remarkably consistent with the only other standardized experiment of comparable scale, which found that attack rates on model caterpillars also increased toward low latitudes and—though not tested systematically—elevations, also driven by invertebrates (9). Together these experiments suggest that invertebrates play an outsized role in the community dynamics and evolutionary trajectories of tropical and lowland ecosystems at multiple trophic scales (8, 9, 22).

Elevational gradients in interaction strength were either independent of latitude or stronger toward higher latitudes. Shallower gradients in the tropics could be because seed predation saturates at lowland tropical sites (5 of 6 sites <1000 m and <20°N had >90% seed predation, Fig. S1), or because the ecological factors that determine seed predation change more slowly up tropical vs. temperate mountains (38). Shallower elevation gradients in tropical interactions contrast a well-known biogeographic paradigm that mountains are ecologically higher and steeper in the tropics (39, 40). Thus, faster community turnover up tropical vs. temperate mountains need not result in greater changes in interaction intensity.

Our findings suggest testable hypotheses about the biogeographic importance of species interactions. Theory predicts that stronger interactions will produce stronger selection (28), so our finding of higher seed predation in the tropics supports a greater evolutionary role for interactions among tropical species (7, 19, 22). More intense interactions toward low elevations and in forest vs. alpine habitat suggests stronger biotically-mediated selection in these environments as well, comparisons that have received less attention than latitudinal contrasts. Plant-herbivore interactions are proposed to drive the impressive diversity of tropical leaf-eating insects and leaf defenses (22). Yet, seed predation affects plant fitness more directly than herbivory, so should impose even stronger selection and demographic effects (21). If defenses cannot fully compensate for increased predation, our results predict greater seed limitation of plant demography and migration rates (26) in high-predation (low-latitude, low-elevation, forested, and less-seasonal) ecosystems.

Geographic patterns in interaction strength have mostly been discussed in terms of their effects, rather than their causes. Seed predation intensity differed both between and within biomes, suggesting both discrete and continuous ecological causes. Temperature seasonality has previously been associated with seed predation intensity, but the direction of the association is inconsistent (negative in this study, positive in 24, 37). The next step in understanding biogeographic patterns in interaction intensity will be to combine large scale, standardized experiments with tests of potential mechanisms and fine scale measurements of underlying gradients (e.g. 37). Identifying general mechanisms underlying interaction gradients would fundamentally improve our understanding of large-scale feedbacks between abiotic gradients and biotic communities (23). Generalizable relationships provide a mechanistic basis for predicting the relative ecological and evolutionary importance of biotic interactions in shaping species distributions, diversity, ecological networks, and resulting responses to of ecological communities to global change (41, 42).

## Acknowledgments

We thank Chris Muir, Victor Vásquez Reyes, Scott Tremor, Carlos García-Jiménez, Lana Bolin, John Godlee, Sandra Angers-Blondin, Javier Tolome, Pablo Sierra, Haley Branch, Joe Boyle, Santeri Lehtonen, Cameron Cosgrove, Mathew Little, Lee Dyer, Hector Barios, Jakob Assmann, and Segundo Chimbolema for help with fieldwork, Adam Davis and Lee Dyer for help with structural equation modelling, and Greg Langston for help with mapping. Funding: UBC Biodiversity Research Grant (ALH), INECOL (proy. 20030-10796 to KM), NSF (grant DEB-1442103 to L. Dyer), and the San Diego Natural History Museum (to S. Tremor). Author contributions: Authorship is in order of contribution. ALH designed and coordinated the study, analyzed the data and wrote the manuscript, all authors contributed to data generation and editing. Authors have no competing interests. All data and code will be archived at the Dryad digital repository. Special thanks to Gustavo Suarez of Fundación Colibrí, who passed away in 2017, for helping us establish sites in La Mesenia-Paramillo Reserve, Colombia.

## Materials and Methods

### Field experiment

We ran this distributed experiment from 2015-2017 to test for latitudinal and elevational gradients in biotic interactions, namely post-dispersal seed predation. Collaborators were recruited who were already working on elevational gradients so that the experiment could be easily integrated into existing field work and permits. Each collaborator established a transect of 4 to 5 sites that spanned at least 1000 m of elevation, or as much elevation as possible given the terrain (Table S1). Site locations were occasionally adjusted between runs; in these cases, a transect consisted of four sites during each experimental run but had 5 to 6 different sites in total (Table S1, Fig. S1).

To increase our ability to detect a latitudinal gradient, if one existed, we standardized our sites, timing, methods and seeds as much as possible. All sites were on the continental Americas, within ~300 km of the Pacific coast (Fig. 1). This latitudinal gradient includes many protected areas along an essentially unbroken mountain chain, minimizing differences in seed-predator communities due to large-scale dispersal barriers (e.g. oceans). Sites were in natural areas, though most had a history of light human disturbance (e.g. logging >100 years ago, nearby park roads and/or hiking trails to enable access).

Experiments were conducted when at least some plant species were dispersing seed at each site. We could not standardize site phenology more precisely, because tropical and temperate sites differ in seasonality, and because growing season phenology differs among elevations at the same latitude on any given date. To better capture average seed predation intensity at each site, the experiment was conducted multiple times at most (15 of 18) transects (median replicates per site = 4). Replicates were separated by at least two weeks and usually several months, between 2015 and 2017.

We used agricultural seed species to ensure seeds were not local to any site, breaking potential coevolved or learned associations between seeds and seed predators. We bulk-purchased organic seeds from the same supplier throughout the experiment: sunflower seeds from Community Natural Foods and oat seeds from West Coast seeds, both in British Columbia, Canada. Seeds were heat sterilized in drying ovens at the University of British Colombia to ensure they would not germinate if dispersed intact by seed predators. Seeds were heated at 110 °C for 1 hr (a slight modification of (24)). Because of this relatively low temperature, seeds did not change noticeably in colour or smell. Sterilized seeds were mailed to collaborators and stored in a cool place in odour-proof containers (e.g. clean glass jars) until use. We used only intact seeds, so that any damage to the seed was unambiguously due to seed predators.

Collaborators used a consistent protocol to quantify seed predation intensity. For each run of the experiment (one date at one transect), we set out 30 seed depots per site: 10 depots of five oat seeds and 20 depots of eight sunflower seeds. Depots alternated between species (2 sunflower depots, 1 oat, repeat). Depots were placed at least 5 m from any walking trail, and at least 5 m from each other in 2015 and ≥10 m apart from 2016 onward – otherwise protocols did not change among years. Seeds were placed directly on the ground in a shallow depression (0.5-3 cm deep and 5-10 cm diameter), a natural depression if available or made by the experimenter.

To facilitate refinding seeds, we removed litter (if any) from the depot so that seeds were set on bare ground. Care was taken not to disturb/remove vegetation outside of the 5-10 cm depot area. Depots were marked with a popsicle stick at the depot edge and green flagging tape 1-2 m from the depot (Fig. S2).

During ca. half of the runs (25 of 56 runs, 14 of 18 transects), we caged some depots to assess invertebrate-only seed predation. We only caged sunflower seeds as oat seeds rarely showed signs of invertebrate predation. We excluded vertebrates from 3 to 4 sunflower depots per site, spaced evenly across the 20 sunflower depots. Cages were made of half-inch square (2.54 × 2.54 cm) wire mesh—large enough for many invertebrates to enter, but the same mesh size as pet mouse cages and rodent-exclusion cages in other experiments (37) to prevent vertebrates from entering. Mesh was shaped to form conical cages ca. 12 cm high and 15 cm diameter (Fig. S2C). Cages were secured over depots using metal pins. If a cage was compromised (dug under by rodents, pulled off by small mammals, or trampled by large mammals) data from that depot were excluded.

We quantified predation 24 h after depots were set out. Sites along a transect were generally set up in one day and checked 24 h later in the same order, but less accessible transects (Ecuador, 19.6° Mexico, 3.3° Colombia) were split into two groups of sites and the experiment run over >2 days. After 24 h we took a photograph of each depot and seed remnants, if any. We recorded the number of intact seeds, seeds with partial predation (i.e. part of the seed remained), the type of damage, signs of predators (rodent faeces (Fig. S2E), chew marks on popsicle stick markers, slug or snail slime trails), or actual predators seen eating seeds (usually slugs or ants, Fig. S2D). All materials were then removed.

### Data manipulation

We calculated seed predation as: proportion of seeds eaten = (seeds partially eaten + seeds missing) / seeds set out (24). This metric combines per-capita effects (seed consumption/removal per granivore) and population size effects, capturing their combined population-level effect (28, 43).

The hypothesis that latitudinal and elevational patterns in biotic interactions exist at a level that influences the long-term ecology and evolution of communities is about the overall average level of interaction intensity. Because we are interested in mean seed predation intensity at each site, we calculated the average proportion of predation across depots of each seed type (sunflower vs. oats) and each predator-exclusion treatment (open = all seed predators vs. caged = invertebrate predators only) at each site, for each date the experiment was run. This generated 1 to 6 measures of mean seed predation intensity per seed type per exclusion treatment per site, depending on how many times the experiment was run per site (1 to 6). Averaging across depots also accounts for non-independence of depots within a site on a given date (i.e. removes pseudo-replication at the depot level), and made the data conform to a binomial distribution by eliminating overdispersion. Models using averaged data converged better (no warnings) than models that used the raw, unaveraged data and incorporated extra random effects to deal with nonindependence of depots and overdispersion (see *Statistical analyses* below).

### Environmental data

Climate, productivity, and biodiversity have all been invoked to increase the strength of biotic interactions toward the tropics. We compared their relative ability to explain spatial variation in seed predation using data from gridded databases extracted for each of our 79 site locations. Following Orrock *et al*, who found significant relationships between abiotic variables and predation on oat seeds in grasslands (24), we tested climate variables from WorldClim (temperature and precipitation mean and variation (44)), and actual evapotranspiration (AET) as a measure of ecosystem productivity (NASA (45)). We extracted diversity data from Biodiversitymapping.org, which has compiled all vertebrate range maps produced by the IUCN and BirdLife International (46, 47). For predation on sunflower seeds, we used total vertebrate species richness as a heuristic for total seed predator diversity, because vertebrate ranges are well mapped and richness data readily available. This is clearly a rough measure, as many vertebrates do not consume seeds and many seed predators are not vertebrates. However, for our study area—Pacific coast of the Americas—vertebrate richness varies similarly to country/state-level species richness for ants, a major invertebrate seed predator, so we think it provides a useful heuristic until better data are available. For oat seeds, which are predominantly eaten by small mammals, we used species richness of rodents and shrews, the primary small mammal seed consumers (48).

### Statistical analyses – Latitudinal and elevational patterns

We used generalized linear mixed models (49, 50) to quantify the effect of latitude, elevation, seed type, and their interactions on seed predation. Because the seed predation data were proportional, we used a binomial error distribution and a logit-link function. As sites on a transect may vary together temporally in seed predation intensity (e.g. with regional pest outbreaks), and repeated measures of a single site are not independent, all models include a random intercept for the date of each experiment and a random intercept for each site. We also tried analyzing the data including the individual seed depots as the base-level of the hierarchy and additional random factors for ‘sitexdate’ to account for non-independence of depots at a given site on a given date, and ‘depot’—an individual-level random factor to resolve overdispersion (50). These models gave the same qualitative results (i.e. included the same fixed effects in the final model, yielded the same rank of factors within fixed effects, and had the same slope direction for continuous factors). However, they converged poorly (i.e. produced multiple convergence warnings), so we chose to present simpler models using data averaged at the depot-level of the experimental design.

We ran one model per hypothesis with the following fixed factors (initial models included all possible interactions). M1) We tested whether total seed predation differed between temperate and tropical zones, including transect latitude as a categorical variable (>23.5 °N = temperate, <23.5 °N = tropical), elevation, and seed type. Categorical latitude is consistent with biogeographic hypotheses that compare ‘the tropics’ to ‘the temperate zone’, without necessarily invoking a continuous gradient (17). All other models consider continuous latitude. M2) We tested whether total seed predation declined continuously with increasing latitude and elevation, including transect latitude (decimal degrees), elevation (masl), and seed type. M3) We tested whether geographic patterns in invertebrate-only predation differed from patterns in total predation. This model considers only sunflower seeds, as we did not exclude vertebrates from oat seeds. This model included only sites and dates where invertebrate predation was measured. Factors were latitude, elevation, and exclusion treatment (all predators vs. invertebrates only).

Models initially included all possible interactions among factors. We assessed interaction significance using sequential likelihood ratio tests comparing models with and without the interaction using a χ^2^ distribution (57). Non-significant interactions (a = 0.05) were dropped from models, as model simplification improved convergence of mixed models (i.e. eliminated convergence warnings). Seed type was always significant—sunflower seeds were more heavily predated than oat seeds—but never interacted with other main effects. Results are shown in Table S2. The main paper presents results for sunflower seeds; comparable results for oat seeds are presented in Fig. S3.

### Statistical analyses – Effect of biome (categorical mechanism)

To test whether latitudinal and elevational patterns were explained by differences in predation among biomes, we classified each site relative to local treelines: above upper tree line (alpine, tundra, paramao), below lower tree line (grassland, desert), or between tree lines (forest).

Treeline elevations were determined by coauthors if the transect was close to tree line.

Otherwise, the upper tree line was determined using Google Earth and the nearest mountain high enough to reach tree line (Table S1). **M4**) We first tested whether total seed predation differed among biomes, including latitude, elevation, and seed type as additional predictors (full model: seed predation ~ biome * latitude * elevation * seed.type + (1|sitelD) + (1|date)). Model reduction was as above, and biome estimates were extracted from the reduced model using least squared means (lsmeans command, lsmeans package (52)). While we included latitude and elevation in the model to account for their effects, we did not use this model to test for latitudinal and elevational effects within biomes, as we did not have even elevational and latitudinal coverage above treeline (only two tropical sites, both from the same transect) or below treeline (sites covered a narrow range of latitudes and elevations; Fig. S1). Instead, we ran separate models (**M5**) testing for latitudinal and elevational patterns in total and invertebrate seed predation in forested sites, for which we had good geographic coverage (Fig. S1). Statistical results are reported Table S2.

### Statistical analyses – Continuous mechanisms (climate, productivity, biodiversity)

We tested whether seed predation was correlated with climate, productivity, and biodiversity.

The variables we tested were motivated by the *a priori* hypotheses outlined in the main paper, and a previous large-scale analysis of environmental effects on seed predation (24). <u>Climate</u>: We tested the climate parameters used in Orrock *et al’s* analysis of climate vs. oat seed predation in North American grasslands (24): Mean annual temperature, Temperature annual range (max temperature of warmest month – min temperature of coldest month), Annual precipitation, and Precipitation seasonality (the coefficient of variation). Long term averages (1950-2000) were downloaded from the WorldClim website (worldclim.org, accessed 2017 (44)) for each of our 79 sites at the finest available spatial resolution (1×1 km). <u>Productivity</u>: As in (24), we also tested the relationship between predation and Annual actual evapotranspiration (AET), the water entering the atmosphere via plant respiration and evaporation from soils. For each site location we downloaded AET data for 2000-2013 from NASA’s MODIS Land Science Team website (modis-land.gsfc.nasa.gov (45). Biodiversity: As a proxy for absolute biodiversity, we downloaded the best available diversity data: total vertebrate species richness for 10 × 10 km grid cells (BiodiversityMapping.org (47)). Equivalent data for invertebrates are not available, so we assume that diversity of seed predators correlates with species richness of vertebrates (sunflower seeds), of rodents and shrews (Rodentia and Eulipotyphla; oat seeds).

We tested for correlations among potential explanatory variables (climate variables, AET, species richness), latitude, and elevation. For both the entire data set and the subset of sites at which the vertebrate-exclusion treatment was added, latitude was significantly correlated with most variables, which were also generally correlated with each other (Fig. S3). Elevation and rodent+shrew richness showed the fewest significant correlations with other variables (Fig. S3). We used structural equation modelling to test the mechanistic relationships among correlated predictor variables (53, 54).

Additional manipulations made data suitable for structural equation modelling. First, to deal with repeated measures of individual sites we averaged the data a second time to get one data point per seed type per caging treatment per site. Second, we arcsin transformed data to make it normally distributed. This yielded 79 data points for total predation on sunflower seeds, n=79 for total predation on oat seeds, n=60 for invertebrate predation on sunflower seeds. We analysed these three seed × predator types independently, to allow for varying biogeographic effects on consumption by different predator guilds (8, 9). Finally, to improve model fits we standardized the response and predictors in each data set to mean=0 and SD=1 (54).

We first made a conceptual model, which was too complex to test with the collected data, but clarified our understanding (and assumptions) about how predictors could affect each other and seed predation (Fig. S4). Latitude and elevation are exogenous variables, whose values do not rely on values of other modelled variables. We divided the climate variables into a) a latent variable ‘Climate’ comprised of Temperature annual range, Annual precipitation, and Precipitation seasonality, and b) Mean annual temperature, which we included separately to test our hypothesis that temperature directly affects seed predation via metabolic activity. Both Climate and Mean annual temperature are directly affected by latitude and elevation and directly affect productivity (AET)(55), and Mean annual temperature also directly affects seed predation (Fig. S4). We assumed elevation’s effect on AET was captured by its effect on climate and temperature, but that latitude could affect AET directly via irradiance (solar energy), which increases toward the equator (2, 56). Although productivity is positively correlated with species richness (Fig. S4), global analyses suggest high productivity does not cause high richness (57-59), so we modelled both variables as affected by climate but independent of each other. We let latitude affect species richness, as recolonization of high latitudes post glaciation has resulted in widespread migration lags (60, 61), which should reduce diversity at higher latitudes independently from modern climate (59). While high elevations were also glaciated, the shorter distances required to cross elevational gradients make migration lags negligible (60, 61). Higher seed predator populations could arise from more productive ecosystems (more food available) or more diverse predator assemblages (‘species packing’), so for this reason we modelled direct effects of AET and species richness on seed predation. Finally, to account for effects not captured by other variables, we modelled the direct effect of latitude and on seed predation intensity.

We tested 13 simpler structural equation models (SEMs), which each represented biologically-motivated hypotheses. These SEMs are illustrated in Fig. S4, and the hypotheses they represent are described below. All models include a direct effect of elevation on seed predation, as grid cells for climate, AET and richness data were large enough to encompass multiple elevational sites along steep gradients.

<u>SEM#) SEM name</u>: Hypothesis

<u>SEM1) Climate</u>: seed predation intensity is best explained by climate plus any additional effects of latitude (e.g. on AET or species richness) and elevation.

<u>SEM2) Direct effects</u>: seed predation intensity is best explained by the variables thought to influence it directly.

<u>SEM3) Direct effects no richness</u>: as for SEM2 but excluding species richness, assuming seed predator diversity is either unimportant or poorly captured by vertebrate richness.

<u>SEM4) ‘Orrock’ structured</u>: predation is best explained using the variables that explained seed predation intensity on oat seeds in temperate grasslands in the Americas (24). The effect of latitude is captured by its effects on climate and AET (24). Elevation can affect seed predation indirectly via an effect on AET; this represents the indirect effect of elevation on AET mediated by Mean annual temperature from the conceptual model. We let Annual temperature range affect seed predation directly, to represent the more direct potential effects of temperature vs. precipitation.

<u>SEM5) ‘Orrock’ more linear</u>: as for SEM4 but without the indirect effects of elevation and Annual temperature range via AET.

<u>SEM6) ‘Orrock’ unstructured</u>: as for SEM4 but climate and productivity variables are modelled independently rather than hierarchically.

<u>SEM7) ‘Orrock’ direct only</u>: including only variables identified as important in (24) for which direct effect can be reasonably hypothesized.

Finally, we compared the simplest possible model for each variable thought to have direct effects on seed predation intensity (SEM8-10) or that significantly affected predation in (24) (SEM11-12) to a model with latitude (SEM13). All models also included elevation, with both variables modelled as exogenous with the structure shown in Fig. S4 SEM8.

<u>SEM8) Mean annual temperature + elevation</u>

<u>SEM9) AET + elevation</u>

<u>SEM10) Species richness + elevation</u>

<u>SEM11) Annual temperature range + elevation</u>

<u>SEM12) Annual precipitation + elevation</u>

<u>SEM13) Latitude + elevation</u>

Structural equation models were run using the ‘sem’ function of the R package lavaan (62). We assessed model goodness-of-fit using the Tucker-Lewis Index (<0.9 indicates poor fit, 63) and Root Mean Square Error of Approximation (>0.1 indicates poor fit, 64). Model selection used AIC.

To test whether results were affected by the additional data manipulations required for SEM, we also compared binomial generalized linear models using the main data set (i.e. 1 data point per date per site per seed and caging treatment). We ran one model for each explanatory variable (latitude, AET, species richness or one of the four climate variables) for total seed predation (models included seed type and elevation) and invertebrate predation (models included elevation). Models were compared using AICc.

## Supplementary Text

### Detailed Results of Structural Equation Modelling

Seed predation intensity was best explained by elevation and either Annual temperature range or latitude. Annual temperature range best explained total predation intensity on sunflower seeds, while annual temperature range and latitude equally explained total predation intensity on oat seeds (Table S3). We did not find support for the predicted stronger role of temperature on invertebrate predation, which was best explained by a non-mechanistic model including elevation and latitude (Table S3). Simpler models preformed the best, and additional complexity via indirect effects, latent variables, or more than two predictors increased AIC values and resulted in poor model fits (Table S3). Very similar results were obtained by comparing generalized linear models on the main data set. Total predation was best explained by annual temperature range, elevation and seed type (ΔAIC 8.4 lower than the next best model), whereas invertebrate predation was best explained by a non-mechanistic model including latitude and elevation (ΔAIC 3.1 lower than the next best model). Thus, the results from structural equation modelling are not artefacts of additional data manipulation.

### Author contributions

Authorship is in order of contribution. ALH designed, initiated and coordinated the study. All authors lead at least 6 days of data collection for the project. ALH analyzed the data, and wrote the manuscript. All authors, but especially IMS, contributed to editing the final version.

**Table S1.** Transect details.

| Country | Latitude<br>(decimal °) | Transect elevation (m) |  |  | Tree line elevation (m) |  |
| --- | --- | --- | --- | --- | --- | --- |
|  |  | Min | Max | Total | Upper | Lower |
| USA | 63.5 | 605 | 1430 | 825 | 640 | – |
| Canada† | 61.2 | 1050 | 1835 | 1000* | 1200 | – |
| Canada | 61.2 | 1360 | 2050 | 1000* | 1200 | – |
| Canada | 54.8 | 470 | 1910 | 1440 | 1650 | – |
| Canada | 49.4 | 80 | 1180 | 1100 | 1350 | – |
| USA | 45.4 | 695 | 1875 | 1180 | 1850 | 630 |
| USA | 39.9 | 60 | 2000 | 1940 | 2200 | 100 |
| USA | 35.7 | 930 | 1940 | 1010 | 3100 | 1550 |
| USA | 33.0 | 210 | 1660 | 1450 | 3100 | 1350 |
| Mexico | 31.0 | 520 | 2430 | 1910 | 3000 | 1500 |
| Mexico | 22.1 | 1780 | 2560 | 780 | 2600 | 1800 |
| Mexico | 19.6 | 1325 | 2530 | 1205 | 4000 | – |
| Mexico | 17.2 | 1625 | 2265 | 640 | 3600 | – |
| Costa Rica | 10.4 | 25 | 1005 | 980 | 2700 | – |
| Panama† | 8.8 | 380 | 1380 | 1000 | 3100 | – |
| Colombia | 5.5 | 2180 | 2930 | 750 | 3750 | – |
| Colombia† | 3.3 | 1800 | 2770 | 970 | 3700 | – |
| Ecuador | -0.7 | 440 | 4120 | 3680 | 3750 | – |

\* Total elevation combined over the two transects at this latitude (Yukon, Canada)
† Location of one or two sites were adjusted between runs of the experiment. Thus the transect had four sites each run but five (or six, for Panama) sites in total.

**Table S2.** Results from generalized linear mixed models analyzing total and invertebrate seed predation. Model numbers are as in text. Initial models (grey rows) included all possible interactions, but nonsignificant interactions were dropped from final models (white rows), improving model convergence. Significance of factors and interactions are from likelihood ratio tests comparing the final model to a model without the factor or interaction. *P* values for Chi squared test: * < 0.05, ** < 0.01, *** < 0.001.

| Model | Predation,<br>Biomes | Fixed effects | Latitude | Elevation | Seed type | Lat x Elev | Additional<br>factor |
| --- | --- | --- | --- | --- | --- | --- | --- |
| **M1** | Total, All | Initial: categoricalLat x elev x seed.type |  |  |  |  |  |
|  |  | Final: categoricalLat + elev + seed.type | 20.7<sub>df=1</sub> *** | 7.9<sub>df=1</sub> ** | 50.1 *** | 2.5<sub>df=1</sub><br>*P* > 0.1 | — |
| **M2** | Total, All | Initial: lat x elev x seed.type |  |  |  |  |  |
|  |  | Final: lat + elev + seed.type | 17.7<sub>df=1</sub> *** | 6.9<sub>df=1</sub> ** | 54.7 *** | 1.5<sub>df=1</sub><br>*P* > 0.2 | — |
| **M3**<sup>1</sup> | Total vs<br>Invert, All | Initial: lat x elev x cage.treat |  |  |  |  |  |
|  |  | Final: lat x elev + cage.treat | 39.0<sub>df=2</sub> *** | 21.8<sub>df=2</sub> *** | — | 7.5<sub>df=1</sub> ** | Cage treat:<br>13.7<sub>df=1</sub> *** |
| **M4** | Total, All | Initial: lat x elev x seed.type x biome |  |  |  |  |  |
|  |  | Final: lat + elev + seed.type + biome | 6.5<sub>df=1</sub> * | 2.6<sub>df=1</sub><br>*P* = 0.11 | 54.3 *** | 2.5<sub>df=1</sub><br>*P* > 0.1 | Biome:<br>8.0<sub>df=2</sub> * |
| **M4**<sup>1</sup> | Invert, All | Initial: lat x elev x biome |  |  |  |  |  |
|  |  | Final: lat + elev + biome | 10.0<sub>df=1</sub> ** | 6.3<sub>df=1</sub> * | — | 2.2<sub>df=1</sub><br>*P* > 0.1 | Biome:<br>8.1<sub>df=2</sub> * |
| **M5** | Total,<br>Forest | Initial: lat x elev x seed.type |  |  |  |  |  |
|  |  | Final: lat x elev + seed.type | 8.6<sub>df=2</sub> * | 7.9<sub>df=2</sub> * | 45.2 *** | 4.2<sub>df=1</sub> * | — |
| **M5**<sup>1</sup> | Invert,<br>Forest | Initial: lat x elev |  |  |  |  |  |
|  |  | Final: lat + elev | 10.9<sub>df=1</sub> *** | 5.4<sub>df=1</sub> * | — | 3.7<sub>df=1</sub><br>*P* = 0.056 | — |

<sup>1</sup> sunflower seeds only, including only sites and dates that included the vertebrate-exclusion treatment

**Table S3.** Relative performance of 13 structural equation models (Fig. S4) in explaining the intensity of total and invertebrate-only seed predation. Top models (lowest AIC) are in bold. Model goodness of fit is given by Tucker-Lewis Index (<0.9 indicates poor fit) and Root Mean Square Error of Approximation (>0.1 indicates poor fit)*.

| Structural Equation Model | Total predation<br>sunflower |  | Total predation<br>oat |  | Invert predation<br>sunflower |  |
| --- | --- | --- | --- | --- | --- | --- |
| | ΔAIC | TLI, RMSEA | ΔAIC | TLI, RMSEA | ΔAIC | TLI, RMSEA |
| SEM1: Climate | 475 | 0.46, 0.45 | 471 | 0.44, 0.45 | 361 | 0.56, 0.40 |
| SEM2: Direct effects | 420 | 0.14, 0.61 | 512 | 0.05, 0.51 | 313 | 0.21, 0.58 |
| SEM3: Direct effects no richness | 296 | 0.09, 0.60 | 292 | 0.02, 0.60 | 217 | 0.17, 0.59 |
| SEM4: Orrock structured | 392 | 0.33, 0.51 | 392 | 0.26, 0.52 | 314 | 0.32, 0.51 |
| SEM5: Orrock more linear | 362 | 0.61, 0.38 | 362 | 0.57, 0.40 | 294 | 0.59, 0.39 |
| SEM6: Orrock unstructured | 405 | 0.36, 0.49 | 404 | 0.30, 0.50 | 309 | 0.44, 0.46 |
| SEM7: Orrock direct only | 129 | — | 130 | — | 115 | — |
| SEM8: Mean Temperature | 21 | — | 11 | — | 18 | — |
| SEM9: AET | 16 | — | 7.6 | — | 14 | — |
| SEM10: Species richness | 13 | — | 14 | — | 17 | — |
| SEM11: Temperature range | **0** | — | **0** | — | 11 | — |
|  | (AIC = 638.3) |  | (AIC = 666.4) |  |  |  |
| SEM12: Mean Precipitation | 16 | — | 9.0 | — | 27 | — |
| SEM13: Latitude | 4.1 | — | **1.4** | — | **0** | — |
|  |  |  |  |  | (AIC = 476.4) |  |

\* TLI = 1 and RMSEA = 0 for SEM7 to 13, in which all predictors directly affect the response.

**Table S4.** Summary of top structural equation models explaining predation intensity for each seed × predator type (from Table S3). Data were arcsin transformed and standardized to mean = 0 and SD = 1 before analyses so Estimates are for relative comparison only.

| Predator type,<br>seed type | SEM: independent variables | Estimate (95% CI) | R<sup>2</sup> |
| --- | --- | --- | --- |
| Total predation,<br>sunflower | 11: Annual temperature range<br>Elevation | -0.63 (-0.81, -0.44) ****<br>-0.27 (-0.46, -0.09) ** | 0.366 |
| Total predation,<br>oats | 11: Annual temperature range<br>Elevation | -0.32 (-0.54, -0.10) **<br>-0.14 (-0.36, 0.08) | 0.095 |
|  | 13: Latitude<br>Elevation | -0.21 (-0.44, 0.02) *<br>-0.12 (-0.35, 0.11) | 0.042 |
| Invert predation,<br>sunflower | 13: Latitude<br>Elevation | -0.68 (-0.90, -0.46) ****<br>-0.46 (-0.68, -0.24) **** | 0.390 |

\* $P < 0.05$ , \*\* $P < 0.01$ , \*\*\* $P < 0.001$ , \*\*\*\* $P < 0.0001$ .

**Fig. S1.**
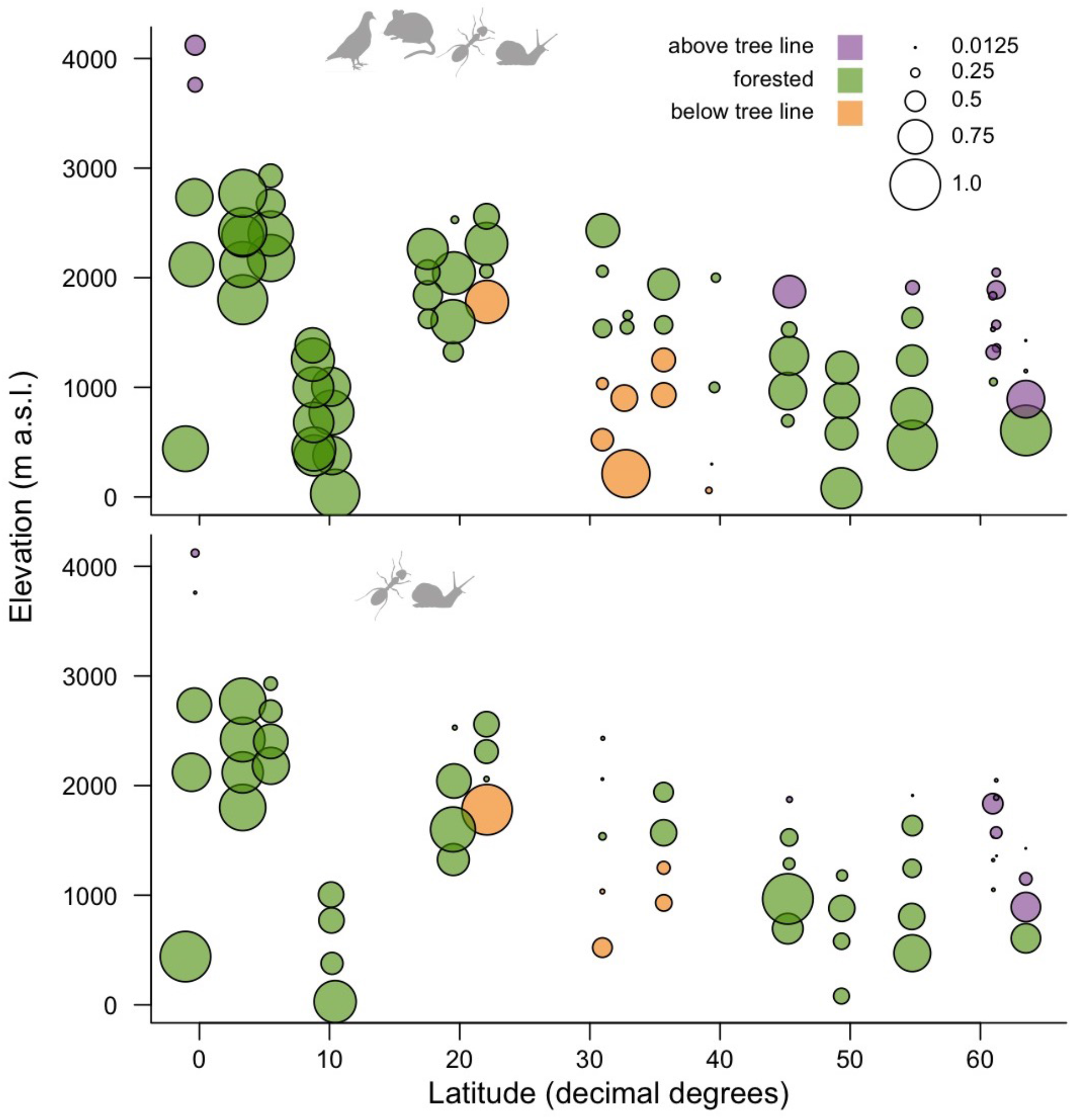
Latitudinal and elevational patterns in mean seed predation. Circle centre shows the latitudinal and elevational location of each site, size shows the mean fraction of predated sunflower seeds, averaged across depots (20/site) and runs (1-6 per site), colour shows site biome. Top panel shows total predation while bottom panel shows predation by invertebrates only (i.e. depots caged to exclude vertebrates).

**Fig. S2.**
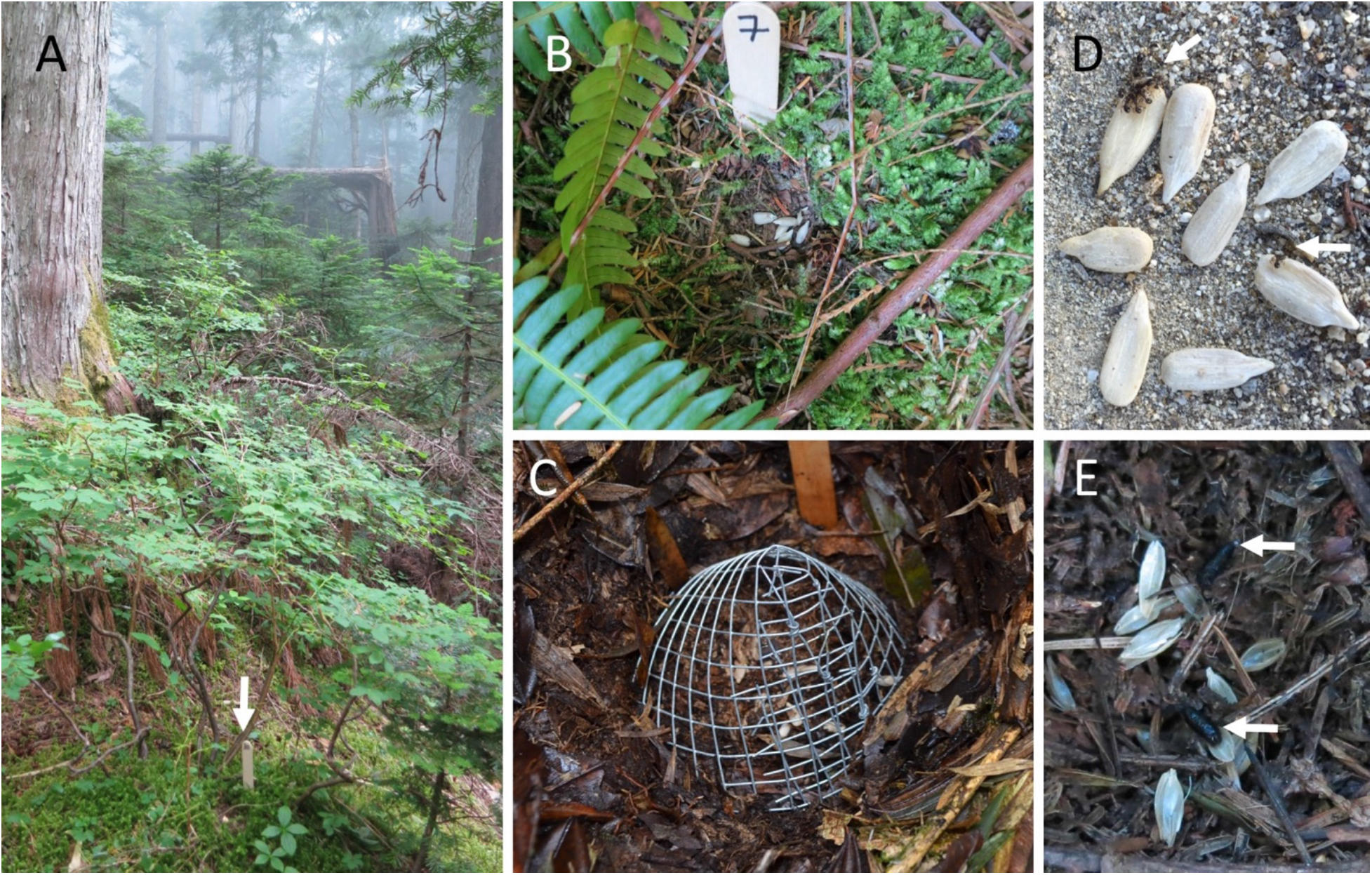
Photos illustrating depot setup at field sites. Depots were not overly visible from >1 m away (A; arrow indicates depot), as care was taken not to disturb the litter and vegetation around the depot (B). Invertebrate seed predation was assessed by excluding vertebrates from some depots of sunflower seeds using ½” mesh cages (C). 24 h after setting out seeds we scored how many were still intact. (D) Sunflower depot with six intact and two partially consumed seeds still being eaten by ants (arrows). (E) Oat depot with no intact seeds—husks peeled from seeds and small mammal droppings (arrows) indicate mammal predation. Photos are from: 49°N in Canada at 80 masl (B), 580 masl (E), and 880 masl (A); 5°N in Colombia at 2120 masl (C); and 31°N in Baja Mexico at 2060 masl (D).

**Fig. S3.**
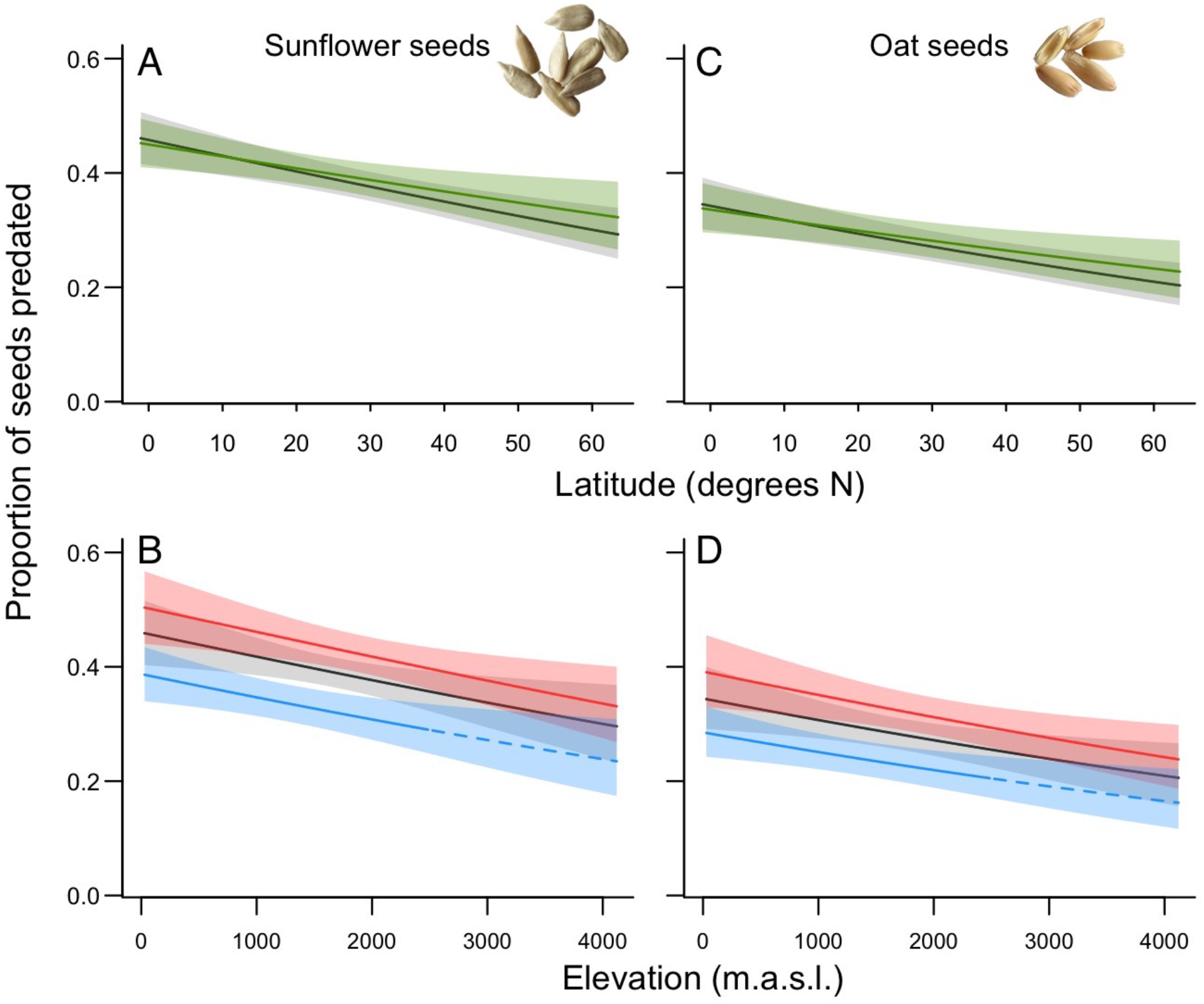
Geographic trends in total seed predation (i.e. vertebrates and invertebrates) on sunflower seeds (A, B) vs. oat seeds (C, D). Lines and shading show trend line +/-95% confidence intervals fitted by generalized linear mixed models (A and B correspond to Fig. 1B and C). All models include seed type as a factor; seed type was always significant, but never interacted with latitude or elevation, i.e. geographic patterns were consistent between seed types. Latitudinal trends (A, C) are shown for the median elevation (1500 m) across biomes (black lines), and in forests (green lines). Elevational trends (B, D) are shown at median latitude (31 °N; black), the tropics (<23.5°N; red), and the temperate zone (>23.5°N; blue). Dashed portion of the line shows trends extrapolated above 2500 m; we had no temperate sites above 2500 m because vegetation stops at lower elevations at higher latitudes.

**Fig. S4.**
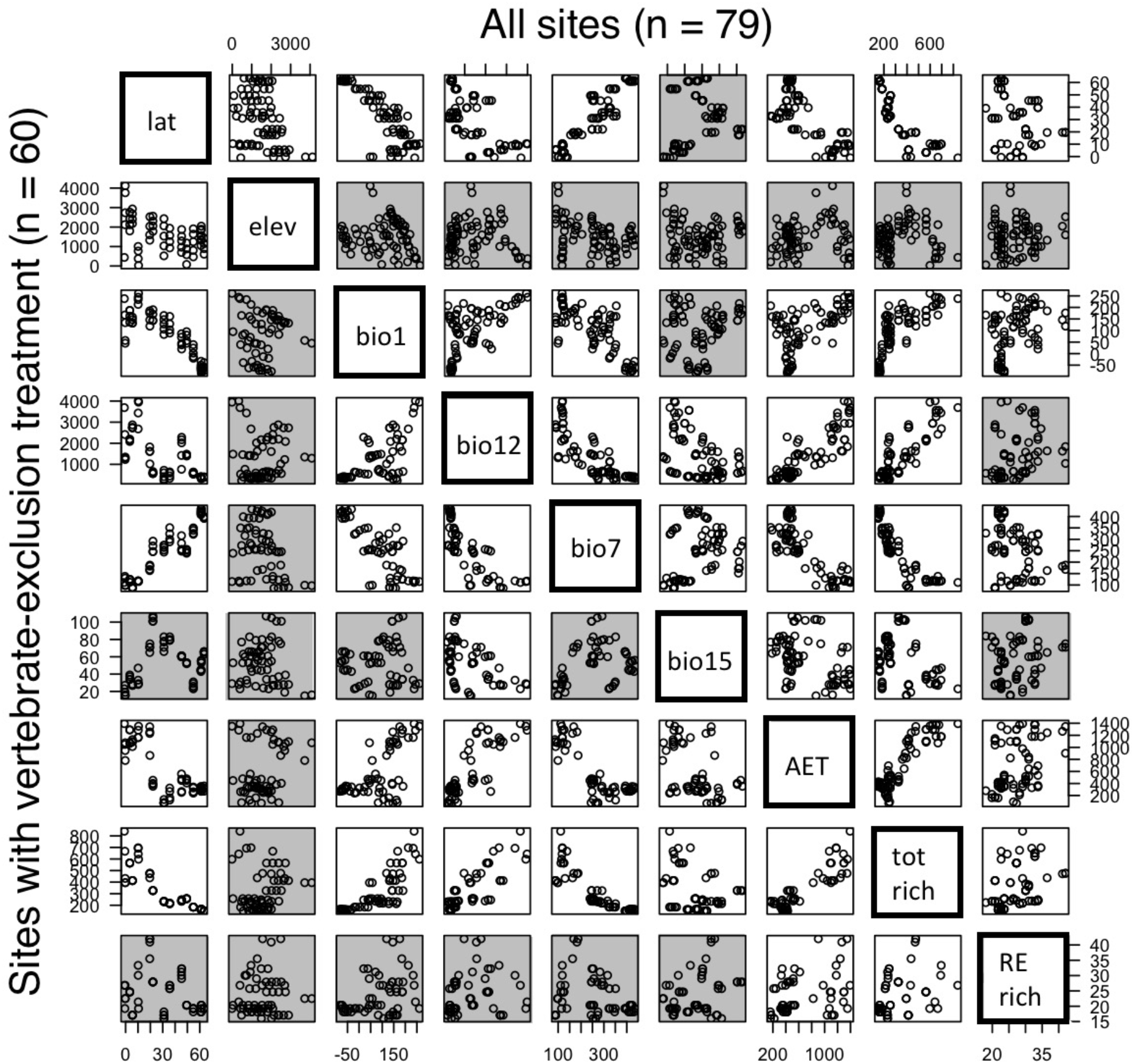
Correlations between continuous variables (latitude, elevation, mean annual temperature (bio1), mean annual precipitation (bio12), annual temperature range (bio7), seasonality of precipitation (bio15), actual annual evapotranspiration (AET), total vertebrate species richness, and rodent and shrew richness. Plots above the diagonal show correlations among all 79 sites, plots below the diagonal show correlations among the 60 sites where the caging experiment was conducted. Grey background indicates correlations that are not significant after correcting for multiple comparisons.

**Fig. S5.**
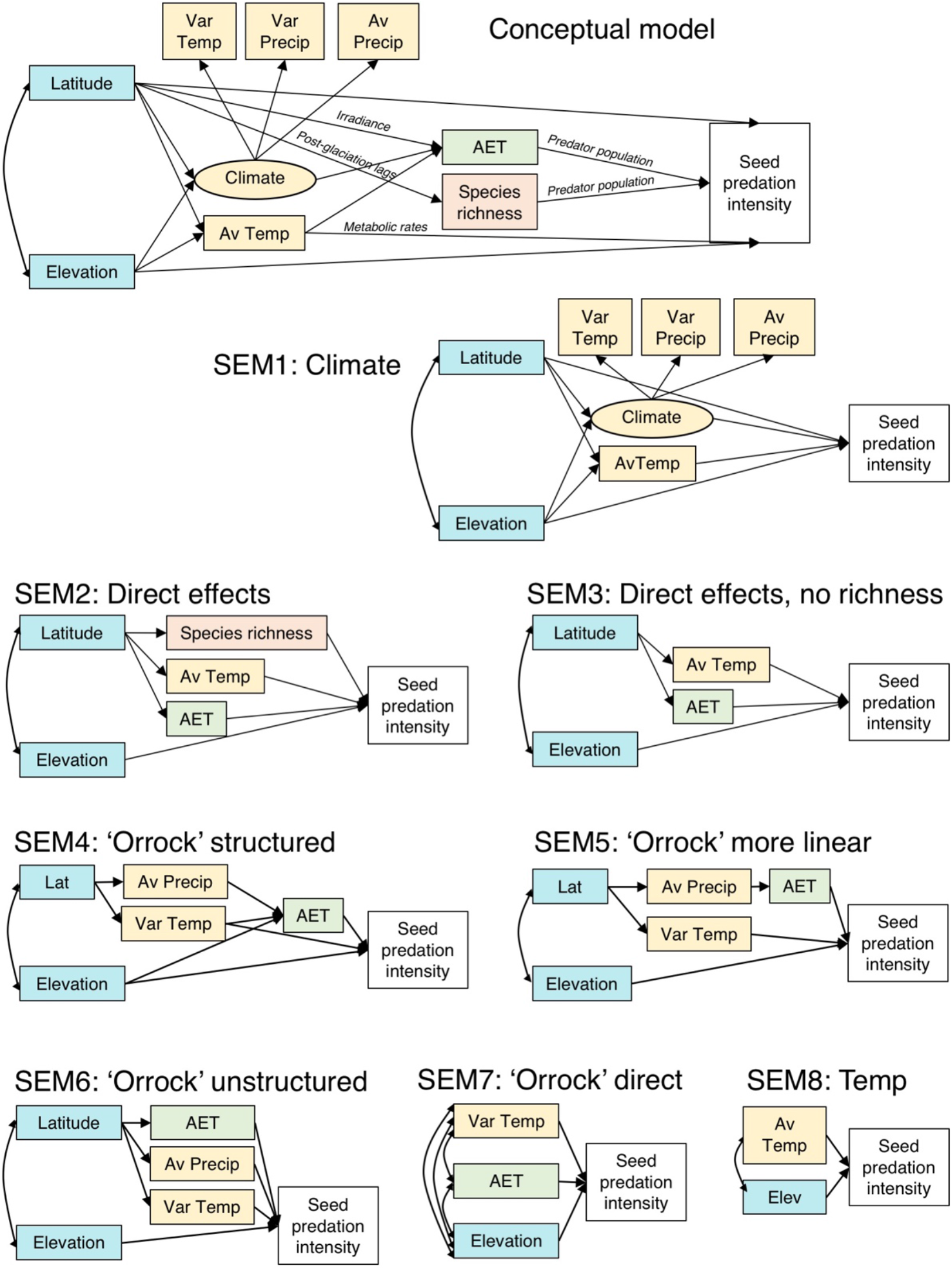
Path diagrams of structural equation models 1 to 8. Manifest variables are boxes, latent variables are ovals, straight arrows denote regression effects and curved double-headed arrows denote correlations. Climate variables (yellow) are Mean annual temperature (Av Temp), Annual temperature range (Var Temp), Annual precipitation (Av Precip), and Precipitation seasonality (Var Precip). Italics along conceptual model arrows give the hypothesized reason for these effects. For ‘Species richness’, models of predation on sunflower seeds use total vertebrate richness, whereas models of predation on oat seeds use the total richness of rodents and shrews.

